# Energetics and proton release in Photosystem II from *Thermosynechococcus elongatus* with a D1 protein encoded by either the *psbA_2_* or *psbA_3_* gene

**DOI:** 10.1101/2023.02.13.528314

**Authors:** Alain Boussac, Julien Sellés, Miwa Sugiura

**Affiliations:** I2BC, UMR CNRS 9198, CEA Saclay, 91191 Gif-sur-Yvette, France; Institut de Biologie Physico-Chimique, UMR CNRS 7141 and Sorbonne Université, 13 rue Pierre et Marie Curie, 75005 Paris, France; Proteo-Science Research Center, and Department of Chemistry, Graduate School of Science and Technology, Ehime University, Bunkyo-cho, Matsuyama, Ehime 790-8577, Japan

## Abstract

In the cyanobacterium *Thermosynechococcus elongatus*, there are three *psbA* genes coding for the Photosystem II (PSII) D1 subunit that interacts with most of the main cofactors involved in the electron transfers. Recently, the 3D crystal structures of both PsbA2-PSII and PsbA3-PSII have been solved [Nakajima et al., J. Biol. Chem. 298 (2022) 102668.]. It was proposed that the loss of one hydrogen bond of Phe_D1_ due to the D1-Y147F exchange in PsbA2-PSII resulted in a more negative *E*_m_ of Phe_D1_ in PsbA2-PSII when compared to PsbA3-PSII. In addition, the loss of two water molecules in the Cl-1 channel was attributed to the D1-P173M substitution in PsbA2-PSII. This exchange, by narrowing the Cl-1 proton channel, could be at the origin of a slowing down of the proton release. Here, we have continued the characterization of PsbA2- PSII by measuring the thermoluminescence from the S_2_Q_A_^-^/DCMU charge recombination and by measuring proton release kinetics using time-resolved absorption changes of the dye bromocresol purple. It was found that *i*) the *E*_m_ of Phe_D1_^−•^/Phe_D1_ was decreased by ∼ 30 mV in PsbA2-PSII when compared to PsbA3-PSII and *ii*) the kinetics of the proton release into the bulk was significantly slowed down in PsbA2-PSII in the S_2_Tyr_Z_^•^ to S_3_Tyr_Z_ and S_3_Tyr_Z_^•^ → (S_3_Tyr_Z_^•^)’ transitions. This slowing down was partially reversed by the PsbA2/M173P mutation and induced by the PsbA3/P173M mutation thus confirming a role of the D1-173 residue in the egress of protons trough the Cl-1 channel.

## Introduction

The water splitting by Photosystem II (PSII) in plants, cyanobacteria, and algae produces the atmospheric dioxygen. PSII consists of 17 trans-membrane proteins and 3 to 5 extrinsic membrane proteins depending on the species [1–4]. The mature PSII binds 35 chlorophylls *a* (Chl-*a*), 2 pheophytins (Phe), 1 membrane b-type cytochrome, 1 extrinsic c-type cytochrome (in cyanobacteria and red algae), 1 non-heme iron, 2 plastoquinones (Q_A_ and Q_B_), the Mn_4_CaO_5_ cluster, 2 Cl^-^, 12 carotenoids and 25 lipids. In the cyanobacterium *Synechocystis* sp. PCC 6803 a 4^th^ extrinsic subunit, PsbQ, has also been found in addition to PsbV, PsbO and PsbU [3].

Among the 35 Chl-*a*, 31 are antenna Chls. After the absorption of a photon by these antenna Chls, the excitation energy is transferred to the photochemical trap that consists of the four Chls; P_D1_, P_D2_, Chl_D1_, Chl_D2_. These 4 Chls, together with the 2 Phe molecules, Phe_D1_ and Phe_D2_, constitute the reaction center of PSII. A few picoseconds after the formation of the excited Chl_D1_*, a charge separation occurs resulting ultimately in the formation of the Chl_D1_^+^Phe_D1_^-^ and then P_D1_^+^Phe_D1_^-^ radical pair states [5,6]. After this charge separation, P_D1_^+^ oxidizes Tyr_Z_, the Tyr161 of the D1 polypeptide, which then is reduced by the Mn_4_CaO_5_ cluster. The electron on Phe_D1_^-^ is then transferred to Q_A_, the primary quinone electron acceptor, and then to Q_B_, the second quinone electron acceptor. Whereas Q_A_ can be only singly reduced under normal conditions, Q_B_ accepts two electrons and two protons before to leave its binding site and to be replaced by an oxidized Q_B_ molecule from the membrane plastoquinone pool, see for example [7–11] for a non-exhaustive list of recent reviews on PSII function.

The Mn_4_CaO_5_ cluster, oxidized by the Tyr_Z_^●^ radical formed after each charge separation, cycles through five redox states denoted S*_n_*, where *n* stands for the number of stored oxidizing equivalents. The S_1_-state is stable in the dark, which makes S_1_ the preponderant state after dark-adaptation. When the S_4_-state is formed, after the 3^rd^ flash of light given on dark-adapted PSII, two water molecules bound to the cluster are oxidized, O_2_ is released and the S_0_-state is reformed, [12,13], see *e.g.* [7-9,14] for recent reviews on the Mn_4_CaO_5_ redox cycle.

The D1 and D2 proteins bear the main cofactors involved in electron transfer reactions. In the *T. elongatus* genome there are three different *psbA* genes encoding a D1 protein [15]. They are tlr1843 (*psbA_1_*), tlr1844 (*psbA_2_*) and tlr1477 (*psbA_3_*). The amino acid sequences deduced from the *psbA* genes, among the 344 amino acids, points a difference of 21 residues between PsbA1 and PsbA3, 31 residues between PsbA1 and PsbA2, and 27 residues between PsbA2 and PsbA3. Whereas the *psbA_1_* and *psbA_2_* genes are contiguous in the genome, with the initial codon of *psbA_2_* located 312 bp downstream of the terminal codon of *psbA_1_* the *psbA_3_* gene is located independently and apart from *psbA_1_* and *psbA_2_*.

The presence of several *psbA* genes is a common feature in cyanobacteria and these genes are, at least partially, differentially expressed depending on the environmental conditions [16,17]. For example, under high light conditions, PsbA3 is predominantly produced [18] whereas the *psbA_2_* transcript is up-regulated under either UV illumination [19] or under micro- aerobic conditions [20]. Some functional differences between PSII with either PsbA1, PsbA2, and PsbA3 have been already characterized and discussed [21–24]. The main findings are briefly summarized below.

The split EPR signal arising from the magnetic interaction between Tyr_Z_^•^ and the cluster in the (S_2_Tyr_Z_^•^)′ state, which is induced by near-infrared illumination at 4.2 K of the S_3_Tyr_Z_ state, is significantly modified, and the slow phases of P_680_^+̣^ reduction by Tyr_Z_ are slowed down from the hundreds of μs time range to the ms time range. These two results [21,24] were interpreted at that time by a change in the geometry of the Tyr_Z_ phenol and its environment, likely the Tyr-O…H…Nε-His bonding, in PsbA2-PSII when compared with PsbA(1/3)-PSII. They also pointed to the dynamics of the proton-coupled electron transfer processes associated with the oxidation of Tyr_Z_ being modified in PsbA2-PSII. From sequence comparison, we proposed that the C144P and P173M substitutions in PsbA2 *versus* PsbA(1/3), respectively located upstream of the α-helix bearing Tyr_Z_ and between the two α-helices bearing Tyr_Z_ and its hydrogen-bonded partner, His-190, were responsible for these changes.

From the observed similar S_1_Tyr_Z_^•^Q_A_^−•^ charge recombination kinetics at 4.2 K in PsbA2- PSII and PsbA3-PSII we predicted that *E*_m_(Q_A_/Q_A_^−•^) in PsbA2-PSII was similar to that in PsbA3-PSII [22].

When the *T. elongatus* genome has only the *psbA_2_* gene for D1, a hemoprotein was found to be present in large amount in cells. This new hemoprotein was identified to be the *tll0287* gene product with a molecular mass close to 19 kDa. The overall structure of Tll0287 was found to be similar to some kinases and sensor proteins with a Per-Arnt-Sim-like domain rather than to other c-type cytochromes. The fifth and sixth axial ligands for the heme are Cys and His which results in a low *E*_m_ (∼ -255 mV *vs* NHE). Possible functions of Tll0287 as a redox sensor under micro-aerobic conditions or a cytochrome subunit of an H_2_S-oxidizing system were proposed taking into account the environmental conditions in which *psbA2* is expressed, as well as phylogenetic analysis, structural, and sequence homologies [23]. Interestingly, when the promotor of the *psbA_2_* gene in a *ΔpsbA_1_, ΔpsbA_3_* deletion mutant is replaced by the promotor of *psbA_3_*, Tll0287 is no longer produced at a level detectable by EPR in whole cells (Sugiura and Boussac, unpublished data) thus suggesting that the promotor of *psbA_2_* has a role in trigerring the expression of Tll0287.

By studying PsbA3/Pro173Met and PsbA2/Met173Pro site-directed mutants we concluded that the Pro173Met substitution in PsbA2-PSII versus PsbA3-PSII was an important structural determinants of the functional differences between PsbA2-PSII and PsbA3-PSII. For example, in PsbA2-PSII and PsbA3/P173M-PSII, we found that the oxidation of Tyr_Z_ by P_680_^+●^ was specifically slowed down during the transition between S-states associated with proton release. We thus proposed that the increase of the electrostatic charge of the Mn_4_CaO_5_ cluster in the S_2_ and S_3_ states could weaken the H-bond interaction between Tyr ^●^ and D1/His190 in PsbA2 versus PsbA3 and/or induce structural modification(s) of the water molecules around Tyr_Z_ [24].

Very recently, the PsbA2-PSII and PsbA3-PSII structures were solved at a 1.9 Å resolution [25]. Based on the longer distance modelled from the crystallographic data between Phe_D1_ and D1-Y126 and the loss of an H-bond due to the D1-Y147F substitution in PsbA2-PSII compared to PsbA3-PSII, a destabilization of Phe_D1_^-^ was expected in PsbA2-PSII [25]. Consequently, the authors in [25] expected a more negative redox potential for Phe_D1_ in PsbA2- PSII than in both PsbA3-PSII and PsbA1-PSII. Part of the increase of the E*m* of Phe_D1_ from - 522 mV in PsbA1-PSII to -505 mV in PsbA3-PSII [26–28] has been explained by a stronger hydrogen bond between Phe_D1_ and Glu130 of D1 in PsbA3-PSII when compared to PsbA1- PSII, *e.g.* [29], with Gln130 in D1.

Finally, the change of D1-Ser270 in PsbA1-PSII and PsbA2-PSII to D1-Ala270 in PsbA3-PSII results in the loss of a H-bond between D1-Ser270 and a sulfoquinovosyl- diacylglycerol molecule near Q_B_. This may result in an easier exchange of bound Q_B_ with a free plastoquinone, hence an enhancement of oxygen evolution in PsbA3-PSII due to a higher Q_B_ exchange efficiency [25] as suggested earlier [21]. The fourth important observation was that two water molecules in the Cl-1 channel were lost in PsbA2-PSII due to the change of D1- P173M, leading to the narrowing of the channel. The authors in [25] concluded that by affecting the proton transfer this may explain the lower efficiency of the S-state transition beyond S_2_ by delaying the P_680_^+^ reduction in the S_2_ and S_3_ states in PsbA2-PSII as observed in [26]. However, the changes in the environment of Tyr_Z_ predicted in [22,24] are likely too small to be detectable in the structures.

In this work, we have estimated the E*m* of Phe_D1_ in PsbA2-PSII from thermoluminescence measurements and the kinetics of the proton release by following the time-resolved absorption change of the dye bromocresol purple. We have also monitored the effect of the D1-P173M exchange in PsbA3PSII and the effect of the D1-M173P exchange in PsbA2-PSII. Fig. 1 shows some of the structural changes identified between PsbA2-PSII and PsbA3-PSII in [25] and discussed in the present work.

**Figure 1:**
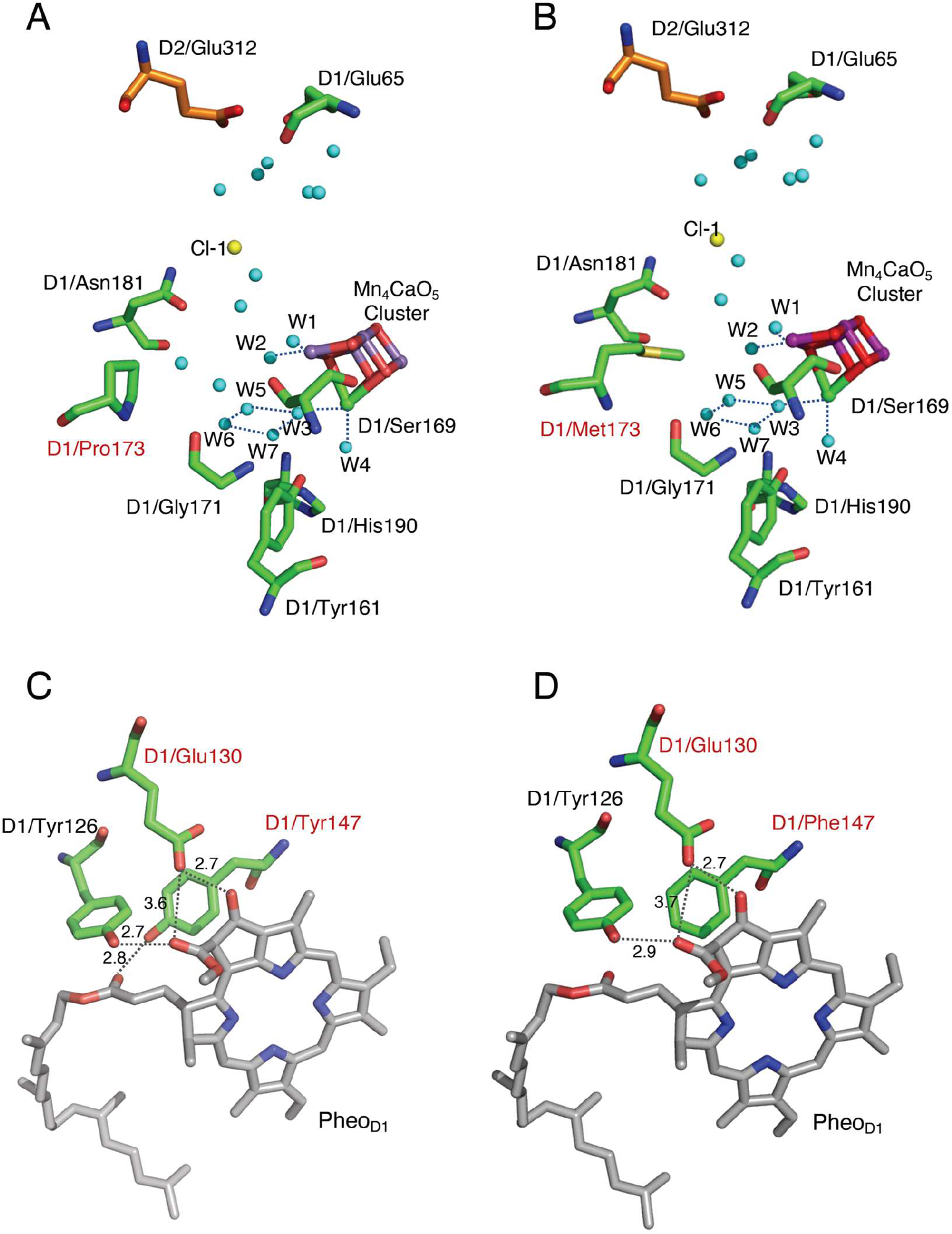
Structural changes between PsbA2-PSII and PsbA3-PSII investigated in the present study. The drawing was done with Mac PyMol 2.5.0 by using the PDB structure 7yq2 for PsbA2-PSII and 7yq7 for PsbA3-PSII [25]. Labels in red characters identify the amino acid residues that differs in PsbA1, PsbA2 and PsbA3.

## Materials and Methods

### PSII samples used

The *T. elongatus* strains used in this study were; *i*) the *ΔpsbA_1_, ΔpsbA_2_* deletion mutant, called WT*3 [30], constructed from the *T. elongatus* 43-H strain that had a His_6_-tag on the carboxy terminus of CP43 [31] and in which the D1 protein is PsbA3; *ii*) the *ΔpsbA_1_, ΔpsbA_3_* deletion mutant, called WT*2 [21], also constructed from the *T. elongatus* 43-H strain, and in which the D1 protein is PsbA2; *iii*) the PsbA3/P173M mutant constructed in WT*3 cells [24]; *iv*) the PsbA2/M173P mutant constructed in WT*2 cells [24]. PSII purification from these 4 strains was achieved as previously described [24]. For the measurements in the presence of bromocresol purple [32], the samples were washed by cycles of dilution in 1 M betaine, 15 mM CaCl_2_, 15 mM MgCl_2_, followed by concentration using Amicon Ultra-15 centrifugal filter units (cut-off 100 kDa) until the estimated residual Mes concentration was ≤ 1 μM in the concentrated PSII samples before the final dilution for the ΔI/I measurements.

### Time-resolved absorption change spectroscopy

Time-resolved absorption changes measurements were done with a lab-built spectrophotometer [33] with the previously described modifications [34]. For the 440 nm- *minus*-424 nm [32,35] the PSII were diluted in a medium containing 1 M betaine, 15 mM CaCl_2_, 15 mM MgCl_2_, 40 mM MES at pH 6.3 (adjusted with NaOH). For the measurements aimed at following the absorption changes of bromocresol purple at 575 nm the samples were diluted in a medium containing 1 M betaine, 15 mM CaCl_2_, 15 mM MgCl_2_, and 150 µM bromocresol purple. PSII samples were dark-adapted for ∼ 1 h at room temperature (20–22°C) before to be diluted at 25 µg Chl/mL then, 100 µM phenyl *p*–benzoquinone (PPBQ) dissolved in dimethyl sulfoxide were added for the two measurements. In addition, for the proton release/uptake measurements, 100 µM ferricyanide from a stock solution, adjusted to pH 6.3 prior to its addition, was also added to avoid contributions from the 2 PPBQ^-^ + 2 H^+^ → PPBQH_2_ + PPBQ reaction. After the ΔI/I measurements, the absorption of each diluted batch of samples was precisely controlled to avoid errors due to the dilution of concentrated samples. The ΔI/I values were then normalized to *A*_673_ = 1.75 [32,35].

### Thermoluminescence measurements

Thermoluminescence (TL) curves were measured with a lab-built apparatus [36,37]. PSII samples were diluted in 1 M betaine, 40 mM MES, 15 mM MgCl_2_, 15 mM CaCl_2_, pH 6.5 and then dark-adapted for at least 1 h at room temperature. Flash illumination was done at - 10°C by using a saturating xenon flash. The constant heating rate was 0.4°C/s. After the dilutions of the PSII samples, and before the dark adaptation, the OD was precisely adjusted to 0.70 at 673 nm (*i.e.* ∼ 10 µg Chl/mL). For the S Q ^-^/DCMU charge recombination, the DCMU (10 µM final concentration) dissolved in ethanol was added before loading the sample into the TL cuvette.

## Results

### Kinetics of the S-state dependent proton release

Fig. 2 shows the kinetics of the absorption changes of bromocresol purple at pH 6.3 with PsbA3-PSII (Panel A), same data as in [38], PsbA2-PSII (Panel B), PsbA3/P173M-PSII (Panel C), and PsbA2/M173P-PSII (Panel D). The measurements were done after the 1^st^ (black points), the 2^nd^ (red points), the 3^rd^ (blue points) and the 4^th^ (green points) laser flashes given on dark-adapted PSII. The analysis of the results in PsbA3-PSII (Fig. 2A) has been previously done in detail [32,38], see also [39] for comparable data in Plant PSII. It is, however, briefly summarized in the next paragraph before to describe the data in the three other samples.

**Figure 2:**
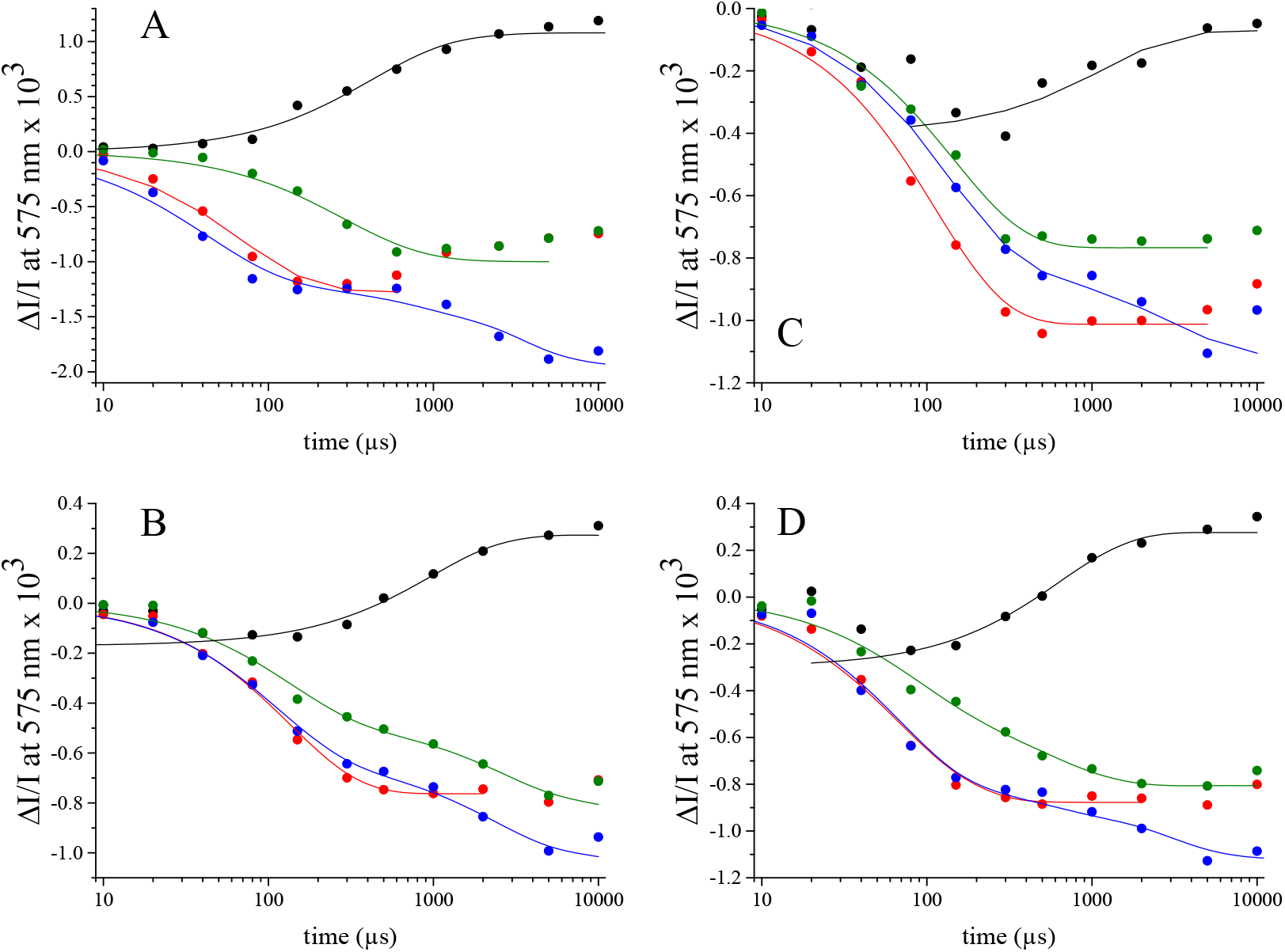
Time-courses of the absorption changes of bromocresol purple at 575 nm at pH 6.3 with PsbA3-PSII (Panel A), replotted from [36], PsbA2-PSII (Panel B), PsbA3/P173M-PSII (Panel C), and PsbA2/M173P-PSII (Panel D). The measurements were done after the 1^st^ (black points), the 2^nd^ (red points), the 3^rd^ (blue points) and the 4^th^ (green points) flashes given on dark-adapted PSII. The continuous lines are the fits of the data using exponential functions. For the 1^st^ flash in PsbA2-PSII, PsbA3/P173M-PSII and PsbA2/M173P-PSII the first data points were not taken into account in the fitting procedure.

In Fig. 2A, the slow increase in the ΔI/I starting at ∼ 100 µs after the flash, and clearly detectable after the first flash in PsbA3-PSII because there is no proton release in the S_1_Tyr ^•^ to S_2_Tyr_Z_ transition, corresponds to the proton uptake with a *t*_1/2_ close to 300 µs (Table 1) occurring *after* the reduction of the oxidized non-heme iron by Q_A_ ^-^ that have a *t* ∼ 50 µs [40].

**Table 1:** t_1/2_ values resulting from the fits of the kinetics in Fig. 2.

| | 1<sup>st</sup> flash | 2<sup>nd</sup> flash<br>($\text{S}_2\text{TyrZ}^\bullet \rightarrow \text{S}_3\text{TyrZ}$) | 3<sup>rd</sup> flash<br>($\text{S}_3\text{TyrZ}^\bullet \rightarrow \text{S}_0\text{TyrZ}$) | 4<sup>th</sup> flash<br>($\text{S}_0\text{TyrZ}^\bullet \rightarrow \text{S}_1\text{TyrZ}$) |
| --- | --- | --- | --- | --- |
| PsbA3 | 317 $\mu\text{s}$ | 48 $\mu\text{s}$ | 30 $\mu\text{s}$ / 1.9 ms | 212 $\mu\text{s}$ |
| PsbA2 | 678 $\mu\text{s}$ | 93 $\mu\text{s}$ | 76 $\mu\text{s}$ / 1.6 ms | 88 $\mu\text{s}$ / 1.9 ms<br>(global $\sim 200 \mu\text{s}$) |
| PsbA3-P173M | 833 $\mu\text{s}$ | 77 $\mu\text{s}$ | 91 $\mu\text{s}$ / 2.1 ms | 101 $\mu\text{s}$ |
| PsbA2-M173P | 440 $\mu\text{s}$ | 50 $\mu\text{s}$ | 47 $\mu\text{s}$ / 1.7 ms | 52 $\mu\text{s}$ / 373 $\mu\text{s}$<br>(global $\sim 100 \mu\text{s}$) |

After the second flash, mainly in the S_2_Tyr_Z_^•^ to S_3_Tyr_Z_ transition, a proton release occurred with a *t*_1/2_ close to ∼ 50 µs. After the third flash, a biphasic kinetics was resolved for the proton release. The fastest phase decayed with a *t*_1/2_ of ∼ 30 µs and the slowest one decayed with *t*_1/2_ ∼ 1.9 ms. These two phases in the proton release corresponds to the two steps in the S_3_Tyr_Z_^•^ → (S_3_Tyr_Z_^•^)’ → S_0_Tyr_Z_ transitions, *e.g.* [39]. After the 4^th^ flash, *i.e.* in the S_0_Tyr_Z_^•^ to S_1_Tyr_Z_ transition, a proton release occurred with a *t*_1/2_ ∼ 200 µs.

For the longest times, a slow drift was observed after all the flashes. It likely includes the end of the proton uptake due to the reduction of the oxidized non-heme iron. This drift does not significantly perturb the interpretation of the kinetics before 1 ms. For example, in this time range, it is clear that the amplitude of the absorption changes are comparable in the S_2_Tyr_Z_^•^ to S_3_Tyr_Z_, S_0_Tyr_Z_^•^ to S_1_Tyr_Z_ and S_3_Tyr_Z_^•^ → (S_3_Tyr_Z_^•^)’ transitions. However, the drift may slightly decrease the amplitude of the slower phase in the S_3_ to S_0_ transition, see [32] for a discussion on this point. In the S_2_Tyr_Z_^•^ → S_3_Tyr_Z_ transition the proton release precedes the oxidation of the Mn_4_CaO_5_ cluster. In the S_0_Tyr_Z_^•^ to S_1_Tyr_Z_ transition the proton release is approximately 4 times slower than the oxidation of the Mn_4_CaO_5_ cluster by Tyr_Z_^•^ [32,39]. A proton release occurs in the first of the two S_3_Tyr_Z_^•^ → (S_3_Tyr ^•^)’ → S_0_Tyr_Z_ transitions whereas the release of the second proton is concomitant to the O_2_ production. The results of the fits are listed in Table 1.

In PsbA2-PSII, PsbA3/P173M-PSII and PsbA2/M173P-PSII (Fig. 2B,2C,2D), the very small decay in the ΔI/I observed after the 1^st^ flash (black points) could be indicative of a proton release in the S_1_Tyr_Z_^•^ to S_2_Tyr_Z_ transition in contrast to the situation in PsbA3-PSII. Alternatively, and most likely, this proton release could occur in a very small fraction (≤ 10 %) of PsbA2-PSII would have been damaged during the washings because the *t*_1/2_ is very close to that measured in Mn-depleted PSII in which the *t*_1/2_ is ∼ 10-20 µs (not shown), *i.e.* in the same time range as the oxidation of Tyr_Z_ by P_680_^+•^ [41].

In PsbA2-PSII, after the second flash (red points), the main decay kinetics had a *t*_1/2_ ∼ 100 µs. Clearly, the proton release in the S_2_Tyr_Z_^•^ → S_3_Tyr_Z_ transition is twice as slow as in the PsbA3-PSII. After the third flash (blue points), the slow phase associated to the O_2_ evolution had a similar *t*_1/2_ (∼ 1.9-1.6 ms) in PsbA3-PSII and PsbA2-PSII. In contrast, the fast phase with a *t*_1/2_ ∼ 80 µs was two to three times slower than in PsbA3-PSII. The situation after the fourth flash (green points) is more delicate to interpret because the miss parameter in PsbA2-PSII is larger than in PsbA3-PSII [21]. Consequently, there is a higher contribution of the S_3_Tyr_Z_^•^ → (S_3_Tyr_Z_^•^)’ → S_0_Tyr_Z_ transitions after the fourth flash. This is particularly clear with the presence of *t*_1/2_ ∼ 1.9 ms phase corresponding to the (S_3_Tyr_Z_^•^)’ → S_0_Tyr_Z_ transition. Nevertheless this complication, the phase with a half time of 200 µs seems to be absent and replaced by a kinetics with a half time of ∼ 90-100 µs. That could suggest that in the S_0_Tyr_Z_^•^ to S_1_Tyr_Z_ transition, the proton release in PsbA2-PSII would be more than twice as fast as in PsbA3-PSII.

Fig. 2C shows the flash dependent kinetics in PsbA3/P173M-PSII. The same question as in the PsbA2-PSII arises for the fast decay phase that has a larger amplitude than in PsbA2- PSII. After the second flash (red points), the proton release with *t*_1/2_ ∼ 80 µs was significantly faster than in PsbA3-PSII and almost similar to that in PsbA2-PSII. After the third flash (blue points), the slow phase for the proton release had *t*_1/2_ close to 2.0 ms, a value close to that in the other samples taking into account the accuracy of the measurement. The fast phase had a *t*_1/2_ ∼ 90 µs, *i.e.* a value comparable to that in PsbA2-PSII and 3 times longer than that in PsbA3-PSII. After the fourth flash (green points) there was no evidence for a large 2.0 ms phase thus indicating a smaller miss parameter than in PsbA2-PSII. Instead, the decay is almost monophasic with a *t*_1/2_ close to 100 µs that is 2 times faster than in PsbA3-PSII and close to the possible fast phase detected in PsbA2-PSII.

Fig. 2D shows the flash dependent kinetics in PsbA2/M173P-PSII and after the first flash (black points) the same comments as in PsbA2-PSII and PsbA3/P173M-PSII can be made. After the second flash (red points), the proton release occurred with *t*_1/2_ ∼ 50 µs that is a value very close to that in PsbA3-PSII and thus twice as fast as in PsbA2-PSII. After the third flash (blue points), the slow phase for the proton release had *t*_1/2_ close to 1.7 ms, a value close to that in the other samples taking into account the accuracy of the measurement. The fast phase had a *t*_1/2_ ∼ 50 µs, *i.e.* a value faster than in PsbA2-PSII and intermediate between PsbA2-PSII and PsbA3-PSII. After the fourth flash (green points), although the global impression seems to suggest an acceleration of the proton release compared to PsbA2-PSII, the complexity of the trace and the number of points necessarily reduced in this kind of experiment does not allow us to conclude.

An additional observation concerns the kinetics for the proton uptake better detectable after the 1^st^ flash. Nevertheless the small number of usable points it is quite clear that this kinetics is twice as slow in PsbA2-PSII than in PsbA3-PSII (Table 1) and that the P173M mutation in PsbA3 slowed down the uptake by a factor ∼ 2 and the M173P mutation in PsbA2 accelerated the uptake also by a factor ∼ 2.

### Time-resolved absorption change differences 440 nm-minus-424 nm during the S-state cycle

The measurement of the 440 nm-*minus*-424 nm difference at various time after a flash is an alternative indirect method to probe the proton and electron transfer kinetics around P_D1_. Indeed, the electrochromic band-shifts in the Soret region of the P_D1_ absorption spectrum at 440 nm takes into account both the proton uptake/release and the electron transfer events, *e.g.* [35,42] and references therein. For the removal of the contributions due to the reduction of Q_A_ the ΔI/I at 424 nm was also measured [42]. A detailed comparison of the measurement using bromocresol purple and the electrochromism 440 nm-*minus*-424 nm in PsbA3-PSII has been done recently [35] and only the main points useful to the present work will be reminded.

Fig. 3 shows the time-resolved absorption change differences 440 nm-*minus*-424 nm in PsbA3-PSII (Panel A) and in PsbA2-PSII (Panel B) after the first four flashes given on dark-adapted PSII. Whereas the kinetics between 5 µs and 10 µs are marred by a small artifactual response of bromocresol purple, discussed previously in [34,43], which makes the 5 µs point more or less unusable, this is not the case here and it can be plotted without any problem.

**Figure 3:**
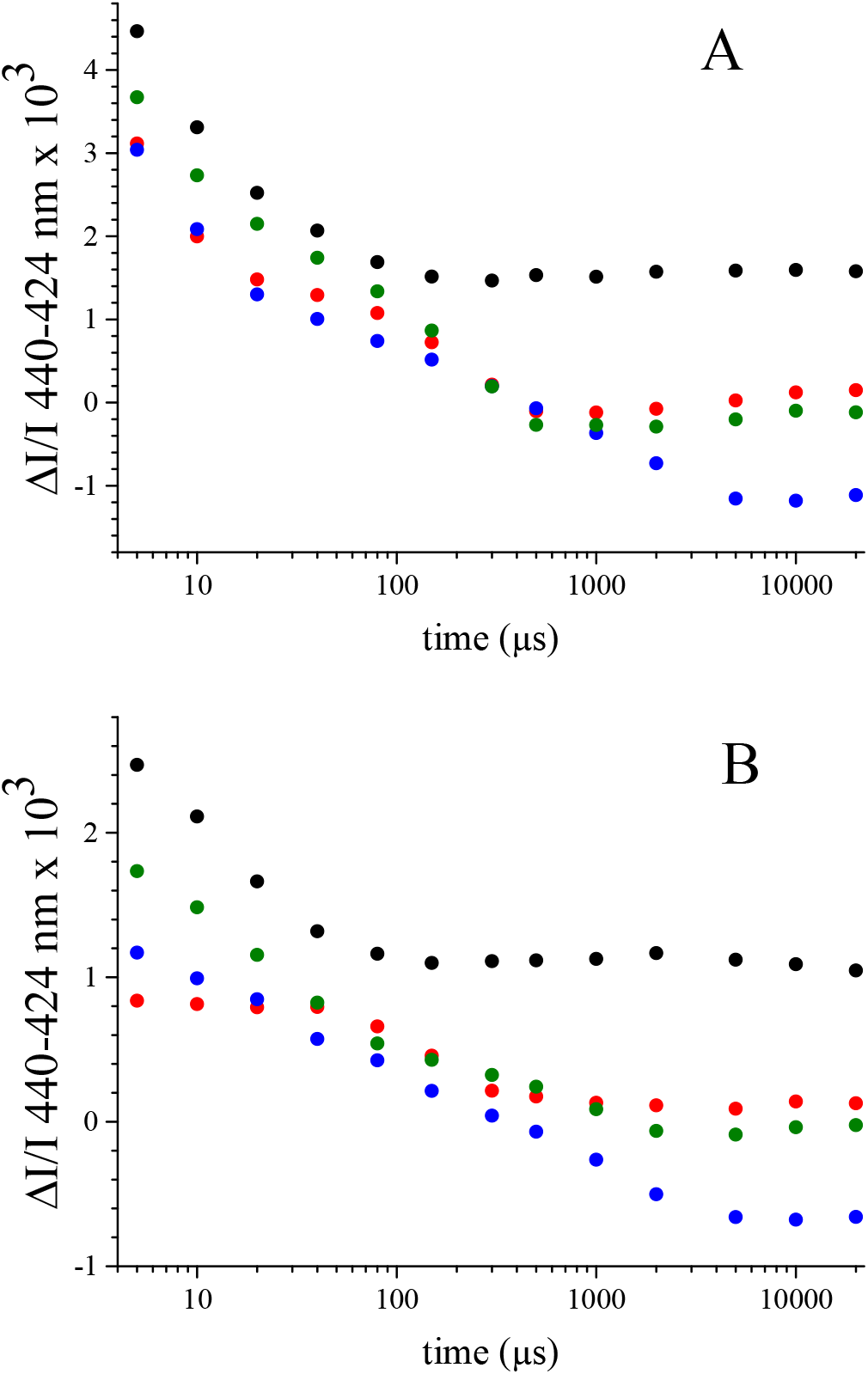
Time-courses of the absorption change differences 440 nm-minus-424 nm after the 1^st^ flash (black), the 2^nd^ flash (red), the 3^rd^ flash (blue), and the 4^th^ flash (green) given to either dark-adapted PsbA3-PSII (Panel A) or PsbA2-PSII (Panel B).

After the first flash (black points) the kinetics corresponds to the reduction of Tyr_Z_^●^ by the Mn_4_CaO_5_ cluster, *i.e.* it corresponds to the S_1_Tyr_Z_^•^ to S_2_Tyr_Z_ transition. This electron transfer reaction had a *t*_1/2_ of ∼ 15 µs in both the PsbA3-PSII (Panel A) and in PsbA2-PSII (Panel B).

After the second flash (red points), the kinetics are dramatically different in the two samples. In PsbA3-PSII (Panel A), the fast phase with a *t*_1/2_ of ∼ 15-20 µs has been identified as a proton movement in the presence of the S_2_Tyr_Z_^•^ state [35], see also [44,45]. This proton movement is likely related, directly or indirectly, to the proton release into the bulk and detected with bromocresol purple with *t*_1/2_ ∼ 50 µs. The slow phase with *t*_1/2_ ∼ 200 µs corresponds to the reduction of Tyr ^•^ in the S Tyr ^•^ to S Tyr transition occurring after the fast proton movement. In PsbA2-PSII (Panel B), the fast phase is missing suggesting that both the proton movement and the electron transfer occur simultaneously with a *t*_1/2_ ∼ 200 µs.

After the third flash (blue points), the kinetics is biphasic in both the PsbA3-PSII and PsbA2-PSII. The slow phase has the same *t*_1/2_ (∼ 1.5 ms) in the two PSII and it corresponds to the (S_3_Tyr ^•^)’ → S_0_Tyr_Z_ transition. In contrast, in PsbA2-PSII the fast phase with *t*_1/2_ ∼ 30 µs is twice as slow as in PsbA3-PSII with *t*_1/2_ ∼ 10-15 µs that is in agreement with the slowing down of the release of the proton into the bulk in this transition in PsbA2-PSII (see above).

After the fourth flash (green points), the kinetics in PsbA3-PSII is also biphasic. By taking into account the kinetics for the release of the proton into the bulk it is clear that the fast phase with *t*_1/2_ ∼ 15-20 µs corresponds to the electron transfer reaction that precedes the proton release which occurs with *t*_1/2_ ∼ 200 µs in the S_0_Tyr ^•^ to S_1_Tyr_Z_ transition. In PsbA2-PSII, the kinetics exhibits 2 phases and possibly 3 phases. The larger miss parameter in this PSII makes the interpretation more difficult. However, if we normalize the amplitude of the two kinetics in the two samples (not shown), the *t*_1/2_ of the fast phase appears similar in PsbA3-PSII and PsbA2- PSII thus suggesting that the electron transfer reaction rates are similar in the two PSII. In contrast, the slow phase seems slower in PsbA2-PSII with *t*_1/2_ ∼ 500 µs. The additional slow phase in PsbA2-PSII with a *t*_1/2_ ∼ 1.5 ms likely corresponds to the (S_3_Tyr ^•^)’ → S_0_Tyr_Z_ transition and this is very likely due to the larger miss parameter.

### Thermoluminescence

Fig. 4 shows the TL signals corresponding to the S_2_Q_A_^-^/DCMU charge recombination in PsbA3-II (black points), PsbA2-PSII (blue points), PsbA3/P173M-PSII (red points) and PsbA2/M173P-PSII (green points). The peak temperature in PsbA3-PSII at ∼ 15°C was as expected in this sample, *e.g.* [34]. In PsbA2-PSII, the peak temperature was shifted by +12°C to ∼ 27°C and the amplitude of the signal was increased by a factor 4.

**Figure 4:**
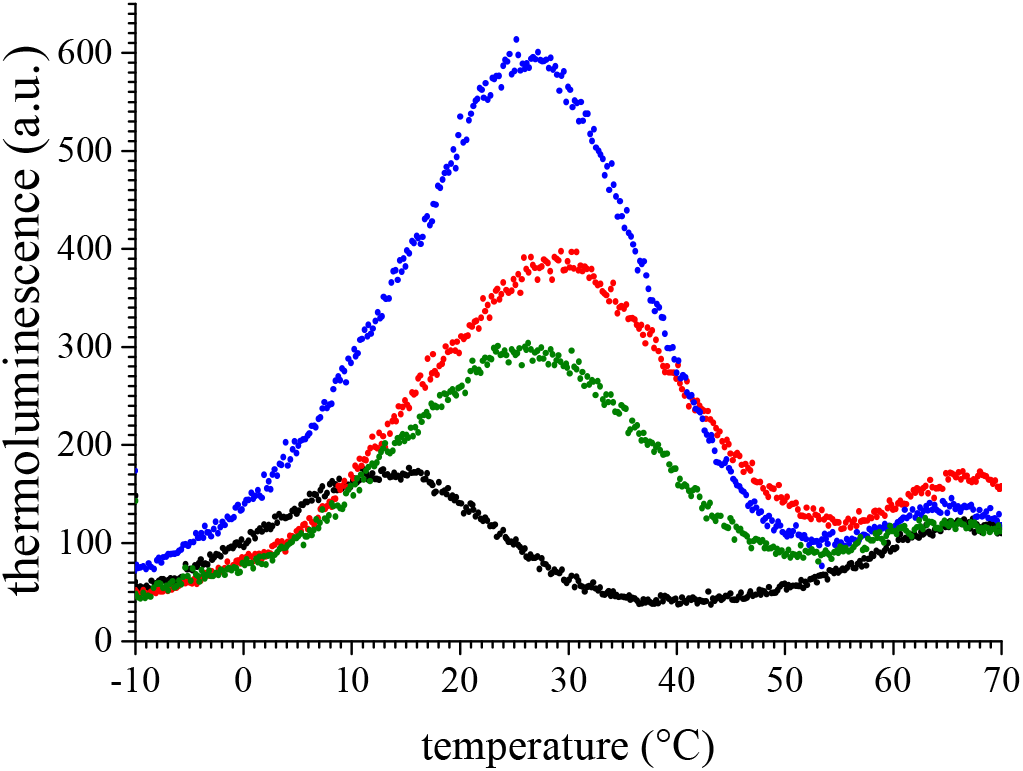
Thermoluminescence curves from S Q ^-^/DCMU charge recombination measured after one flash given at -10°C in PsbA3-PSII (black points), PsbA2-PSII (blue points), PsbA3/P173M-PSII (red points) and PsbA2-M173P-PSII (green points). The heating rate was 0.4°C/s.

In the PsbA3/P173M-PSII the changes in the TL, when compared to the PsbA3-PSII, were qualitatively similar to those observed in PsbA2-PSII. The peak temperature was shifted to 29-30°C but the amplitude was only increased by a factor of ∼ 2 (instead of 4 in PsbA2-PSII). In the PsbA2/M173P-PSII the peak temperature was ∼ 27°C, *i.e.* almost unchanged when compared to PSbA2-PSII. The amplitude was nevertheless reduced by a factor of 2 so that it remained twice that in the PsbA3-PSII.

## Discussion

In previous studies on the properties of PSII depending on the nature of the PsbA [21,22,24] it was shown that in PsbA2-PSII the reduction of P_680_^+•^ by Tyr_Z_ is slower than in PsbA3-PSIIin the S_2_ to S_3_ and S_3_ to S_3_’ transitions, *i.e.* the steps in which a proton transfer/release occurs *before* the electron transfer from the Mn_4_CaO_5/6_ cluster to Tyr_Z_^•^. This slow-down was shown to be significantly either reverted in the PsbA2/M173P mutant and induced in the PsbA3/P173M mutant.

Significant progress has been made recently with the publication of 3D structural models of PsbA2-PSII and PsbA3-PSII with a 1.9 Å resolution [25]. Many computational and experimental studies have addressed the proton channels, *e.g.* [34,46–49], and it is established that the Cl-1 channel is involved in the egress of protons, see *e.g.* [50–53] for some recent experimental evidence. The structure of PsbA2-PSII revealed that with a methione instead of a proline at the position 173 in D1, two water molecules in the Cl-1 channel are lost due to a narrowing of this channel [25]. There is not yet a structure of the PsbA2/M173P-PSII and PsbA3/P173M-PSII, but it seems very likely that this exchange alone can explain the slowdown in PsbA2-PSII of the rates of the proton releases in the S_2_Tyr_Z_^•^ to S_3_Tyr_Z_ and S_3_Tyr_Z_^•^ to (S_3_Tyr ^•^)’ transitions. However, in the (S_3_Tyr ^•^)’ to S_0_Tyr_Z_ transition the kinetics of the proton release is hardly affected thus strongly suggesting that either the egress of the proton in this step occurs *via* a different route that is not kinetically modified in PsbA2-PSII *versus* PsbA3-PSII, or that the removal of this proton from the cluster is the limiting step. In the S_0_Tyr ^•^ to S_1_Tyr_Z_ transition, the data are more difficult to analyse. The most likely reason is the larger miss parameter in PsbA2-PSII and the mutants. However, the proton released in this transition seems to use a different egress pathway, *e.g.* [53]. Presently, it is proposed that this proton initially originates from either protonated O4 [54–56] or protonated O5 [54] or a water molecule bound to the terminal Mn [57], whereas in the S_2_ to S_3_ transition, the first step is a deprotonation of W1 with the proton moving onto Asp61, *e.g.* [57–60]. Computational analysis of the hydrogen bond network in the vicinity of the oxygen evolving complex show that the waters are highly interconnected with similar free energy for hydronium at all locations [48] and this could result in multiphasic kinetics.

On the first flash, in PsbA3-PSII, the *t*_1/2_ of the proton uptake (∼ 320 µs) is ∼ 6 times slower than the electron transfer rate between Q_A_ ^-^ and Fe^2+^ (∼ 55 µs) [40]. Therefore, the slower proton uptake in PsbA2-PSII more likely reflects a change in the proton ingress rather a dramatic change in the electron transfer rate between Q_A_ ^-^ and Fe^3+^, a kinetic rate that nevertheless would remain to determine in PsbA2-PSII. The amino-acids whose p*K*_a_ values have been shown, either experimentally [61,62] or by computational approaches [63], to be tuned by the redox state of the non-heme iron, like the D1-E243, D1-E244, D1-H252 and D1- His215, are conserved in the 3 PsbA variants so that they are not involved in the differences observed between PsbA2-PSII and PsbA3-PSII. The structural changes around Q_B_ in PsbA2- PSII [25] can indirectly affect the kinetics of the proton uptake upon the reduction of the non-heme iron by Q ^-•^. However it is more difficult to explain why the loss of two water molecules in the Cl-1 channel may slow down this kinetics by a factor 2 since the M173P mutation in PsbA3 has the same effect as the PsbA3 to PsbA2 exchange. More experiments are therefore required to determine which amino-acids participate in the ingress/egress of the proton following the reduction/oxidation of the non-heme iron.

Both the magnetic coupling between the Ty_Z_^●^ radical and the Mn_4_CaO_5_ cluster in the S_2_state and the W-band EPR spectrum of Tyr_Z_^●^ in Mn-depleted PSII are modified in PsbA3-PSII, and these modifications have been found induced by the P173M mutation alone in PsbA3-PSII, and reverted by the M173P mutation alone in PsbA2-PSII. This seems to be in disagreement with the 3D structures that reveal no differences in the Tyr_Z_ environment in PsbA2-PSII and PsbA3-PSII. However, *i*) these structures are those with Tyr_Z_ and not with Tyr_Z_^●^, and *ii*) the structural changes causing the differences observed in EPR are probably too small to be detectable in a structure even with a 1.9 Å resolution.

The second important difference between PsbA2-PSII and PsbA3-PSII deduced from the structure, *i.e.* the loss of one H-bond between Phe_D1_ and Y147 in PsbA2-PSII, is an expected lower *E*_m_ of Phe_D1_ in PsbA2-PSII than in PsbA3-PSII [25]. The TL data reported here for the first time in PsbA2-PSII indicate that the energy level of S_2_Q_A_^-^/DCMU is lower in PsbA2-PSII than in PsbA3-PSII and/or the energy level of the P_680_^+•^Phe_D1_^−•^ state is higher in PsbA2-PSII than in PsbA3-PSII [64,65]. The increase in the energy gap between the S_2_Q_A_^-•^/DCMU state and the P_680_^+•^Phe_D1_^−•^ state in PsbA2-PSII when compared to PsbA3-PSII can be estimated to be close to (27-15)/0.4 = 30 meV [64]. We have previously shown that the decay of the S_1_Tyr_Z_^•^ split EPR signal at 4.2 K in PsbA2-PSII was found similar to that in PsbA3-PSII [22]. This suggested that the driving forces for the charge recombination by the non-radiative direct route in the S_1_Tyr_Z_^•^-Q_A_^−•^ state, *i.e.* a route that does not involve the repopulation of the P_680_^+•^Phe_D1_^−•^ state, was similar in PsbA2-PSII and PsbA3-PSII. Since, in addition, in the S_1_ state the Tyr_Z_ oxidation by P_680_^+•^ and the Tyr_Z_^•^ reduction by the Mn_4_CaO_5_ cluster in the S_1_-state remains almost unaffected in PsbA2-PSII, we proposed that the *E*_m_(Q_A_/Q_A_^−•^) value in PsbA2-PSII was similar to that in PsbA3-PSII [22]. Therefore, the differences seen in the TL, *i.e.* an upshift of the peak temperature and a larger amplitude of the TL signal in PsbA2-PSII, are compatible with a lower *E*_m_ of Phe_D1_^−•^/Phe_D1_ as suggested in [25], [64]. Unfortunately, we have no data on the charge recombination at 4.2 K between the S_2_Tyr_Z_^•^ split EPR signal and Q_A_^−•^ in PsbA2-PSII state because this split signal cannot be formed in PsbA2-PSII even at room temperature (likely due to the change in the proton egress discussed above).

The Tyr_Z_ oxidation in the S_2_ state at helium temperatures requires a pre-illumination at 200 K. In a computational study, it has been proposed that “the tautomerization of D1/Asn298 to its imidic acid form enables proton translocation to an adjacent asparagine-rich cavity of water molecules that functions as a proton reservoir and can further participate in proton egress to the lumen” [66]. Since the S_2_Tyr_Z_^●^ state can be formed upon an illumination at 200 K in PsbA3-PSII and PsbA1-PSII and not in PsbA2-PSII, the egress of the proton involved here may occur at 200 K and very likely the proton reservoir is constituted by the water molecules missing in PsbA2-PSII [25]. In addition, since D1/Asn298 is present in the three PsbA, it is not responsible for the loss of the capability of PsbA2-PSII to form the S_2_Tyr ^●^ state.

The TL changes observed in PsbA2-PSII are partially reversed in PsbA2/M173P-PSII and partially induced in PsbA3/P173M-PSII. However, the lack of a symmetrical effect when we compare the data in the PsbA3-PSII *versus* those in the PsbA2/M173P-PSII on one hand and the data in the PsbA2-PSII *versus* those in the PsbA3/P173M-PSII on the other hand indicates that the other amino acid substitutions between PsbA2 and PsbA3 have an effect. Nevertheless, and in conclusion, the data in the present work are in good agreement with the deductions made from the crystal structures of PsbA2-PSII and PsbA3-PSII [26].

Characterizing the functional properties of PSII according to the nature of the D1 protein is the first step in understanding the advantage conferred by each of the D1 isoforms under the conditions where they are preferentially expressed. In PsbA3-PSII, which is up regulated under high light conditions [18], it was found that the less negative *E*_m_ value of the Phe ^−•^/Phe_D1_ couple [18,28] could be responsible for a lower ^1^O_2_ production [28]. Such a change is in agreement with a better resilience of PsbA3-PSII than PsbA1-PSII under high light conditions. However, if the *psbA_2_* transcription is strongly upregulated under micro-aerobic conditions [20], there is so far no quantification of the PsbA proteins in response to this stress and that remains to be done taking into account the possible complex regulation systems already identified in other cyanobacteria [67]. Assuming that PsbA2 is indeed produced under micro- aerobic conditions, a more negative *E*_m_ value of the Phe ^−•^/Phe_D1_ couple in PsbA2-PSII than in PsbA1-PSII and PsbA3-PSII is expected to favor the formation of ^1^O_2_ for all the reasons detailed in [28]. However, in the absence of O_2_ the production of ^1^O_2_ is no longer a problem. The fact that WT*2 cells are well growing in the presence of O_2_ can be explained by the larger energy gap between the Phe Q ^-^ state and the Phe ^-^Q state in PsbA2-PSII than in PsbA1- PSII and PsbA3-PSII, which decreases the thermaly activated repopulation of the Phe_D1_^-^Q_A_ state and therefore favours the direct recombination at the expense of the triplet formation route [68].

## Acknowledgements

This work has been in part supported by (i) the French Infrastructure for Integrated Structural Biology (FRISBI) ANR-10-INBS-05, (ii) the Labex Dynamo (ANR-11-LABX-0011-01) and (iii) the JSPS-KAKENHI Grant in Scientific Research on Innovative Areas JP17H064351 and a JSPS-KAKENHI Grant 21H02447. Bill Rutherford is acknowledged for a careful reading of the manuscript and Yuki Ito and Kosuke Tada for drawing figures.

